# Antiviral Library Screening Leads to Discovery of Novel and Potent Anticlostridial Compounds

**DOI:** 10.1101/2024.08.16.608359

**Authors:** Brice J. Stolz, Ahmed A. Abouelkhair, Mohamed N. Seleem

**Author notes:** Address correspondence to Mohamed N. Seleem, Department of Biomedical Sciences and Pathobiology Virginia-Maryland College of Veterinary Medicine Virginia Polytechnic Institute and State University 1410 Prices Fork Rd, Blacksburg, VA, 24061, USA.

## Abstract

*Clostridium difficile* (*C. difficile*) is a major cause of nosocomial infections, often associated with individuals who have gut dysbiosis from previous antibiotic therapies. *C. difficile* infections have a high recurrence rate and impose significant financial and mortality burdens on the healthcare system. Novel anti-*C. difficile* drugs are urgently needed to treat and reduce the severity and recurrence of infection. In this study, we screened a library of 618 antiviral drugs to identify a potential candidate for repurposing as novel anti-*C. difficile* treatments. Following our preliminary screening, we identified 9 novel compounds that inhibit *C. difficile* at a concentration of 16 µM or lower. Among these, 4 antiviral compounds demonstrated the most potent anti-*C. difficile* activity against a panel of 15 *C. difficile* isolates, with minimum inhibitory concentrations (MICs) comparable to the drug of choice, vancomycin. These include rottlerin (MIC_50_ = 0.25 µg/mL), α-mangostin (MIC_50_ = 1 µg/mL), dryocrassin ABBA (MIC_50_ = 1 µg/mL), and obefazimod (MIC_50_ = 4 µg/mL). All exhibited minimal to no activity against representative members of the gut microbiota. Interestingly, α-mangostin, a natural xanthone derived from the mangosteen fruit, exhibited strong bactericidal action, clearing a high inoculum of *C. difficile* in less than an hour. All other drugs exhibited bacteriostatic activity. Given the characteristics of these compounds, they show great promise as novel treatments for CDI.

## Introduction

Health care-associated infections are one of the greatest burdens on health care systems. Most bacteria responsible for a part of this burden are associated with antimicrobial resistance, with one notable exception: *C. difficile*. Despite lacking a significant antimicrobial resistance profile, *C. difficile* is still considered as an urgent threat and one of the deadliest bacterial infections responsible for approximately 12,800 deaths annually, primarily affecting older populations [1–5]. The incidence of CDI in the United States has increased over the last two decades and has remained relatively stable from 2021 to 2022 with the COVID-19 pandemic seeing a drop in cases due to improved strategies preventing the spread of microorganisms [2, 3, 6]. Typically, the most vulnerable patients receiving antibiotic treatment the target of *C. difficile*. Antibiotic treatment causes gut dysbiosis and, due to a lack of competition in the gut microbiome, promotes the development and colonization of *C. difficile*, which triggers a series of host responses including intestinal tissue necrosis, diarrhea, and pseudomembranous colitis [7–9].

Novel antibiotic therapies approved for *C. difficile* infection (CDI) have been conspicuously lacking. In the past, the recommended drugs for mild to severe CDI were metronidazole and vancomycin, respectively. Fidaxomicin is the most recently approved new antibiotic for CDI, being approved nearly 40 years ago. Due to the high rates of treatment failure and recurrence, metronidazole is no longer recommended; vancomycin and fidaxomicin are now the main therapeutic options for CDI treatments [8, 10, 11]. Despite the potency of these antibiotics against *C. difficile*, recurrence rates can still be significant even if they are an improvement over metronidazole. Fidaxomicin and vancomycin have both seen recurrence rates of 20% or higher with subsequently increasing chances of recurrence after the first episode [12, 13]. There is an immediate and growing need for new and innovative antibiotics to address the shortcomings of existing therapies and their impact on the healthcare system.

Antivirals are one of the most prolific and growing categories of treatments since the first was approved in 1963 [14]. With the advent of COVID-19, antiviral research has become an area of increasing interest and with it many antiviral compounds that are approved for treatment or held back due to their pharmacokinetics [15]. Many antivirals struggle with poor oral bioavailability, something that is necessary in *C. difficile* treatment for it to see significant efficacy towards an infection and would otherwise be overlooked as an antiviral drug.

Screening of antiviral compounds for potential efficacy against *C. difficile* has remained relatively unexplored, despite the need to find novel anti-*C. difficile* drugs and address the postponed discovery of antibiotics for this pathogen. Prior screens and clinical studies have demonstrated the potential of antiviral drugs, such as nitazoxanide, a broad-spectrum antiviral and antiprotozoal drug, against *C. difficile* in comparison to vancomycin [16–18]. Furthermore, antiviral substances have demonstrated potential against other microbes. In a study done on antiviral hypericin, it demonstrated can boost the efficacy of β-lactam antibiotics against methicillin-resistant *Staphylococcus aureus* (MRSA) [19] .

Given the potential of the aforementioned medications, the goal of this work is to screen an antiviral library to find antiviral compounds with comparable potency against *C. difficile*. Screening an antiviral library revealed four antiviral compounds that had strong anticlostridial action activity. These hits were assessed for potency and specificity against a panel of clinical isolates of *C. difficile* and the human gut microbiota, respectively. As well as their mode of action and the time it takes to reach this action when inhibiting *C. difficile*.

## Materials and Methods

### Bacterial strains & reagents

Bacterial strains utilized in this study were sourced from the Centers for Disease Control and Prevention (CDC, Atlanta, GA) Biodefense & Emerging Infections Research Resources Repository (BEI Resources, Manassas, VA) and the American Type Culture Collection (ATCC, Manassas, VA). Phosphate-buffered saline (PBS) (Corning, NY, USA), Brain heart infusion broth (BHI) and De Man—Rogosa—Sharpe broth (MRS) (Becton, Dickson Company, NJ, USA), yeast extract (Fisher Scientific Global Solutions, Suwanee, GA), L-cystine (Thermo Fisher Scientific, Waltham, MA), vitamin K1, resazurin, and hemin (Sigma-Aldrich, St. Louis, MO) were purchased commercially.

### Compounds and libraries

The MCE antivirals library (Cat. No: HY-L027) which includes 618 unique compounds displaying antiviral activity, was purchased from MedChemExpress (Monmouth Junction, NJ). After an initial screening and confirmation, the active hits were purchased commercially as follows rottlerin and dryocrassin ABBA (MedChemExpress; Monmouth Junction, NJ), *α*-mangostin (TargetMol Chemicals; Wellesley Hills, MA), and obefazimod (Ambeed; Arlington, IL). Vancomycin hydrochloride (Gold Biotechnology, Olivette, MO, USA) were included as positive controls.

### Screening assay against *C. difficile*

To discover antivirals with anti-*C. difficile* action, the antivirals library was screened using the microbroth dilution technique, as previously described [20–23], against *C. difficile* ATCC BAA 1870 at a fixed concentration of 16 µM. In brief, the bacteria streaked and grown anaerobically at 37°C for 48 hours on brain heart infusion supplemented (BHIS) agar plates. The antiviral compounds were then added to a 0.5 McFarland solution of *C. difficile* at a concentration of 16 µM. The drugs were then diluted in BHIS broth (∼5 x 10^5^ CFU/ml) and placed in 96-well plates. The plates were then incubated anaerobically at 37°C for 48 hours. Using a BioTek Synergy H1 Microplate Reader with BioTek Gen 5 Microplate Reader and Imager Software, the OD600 was determined. The compounds that inhibited 90% of the bacterial growth in the dimethyl sulfoxide (DMSO) negative control were recognized as potential hits of interest and purchased commercially for further screening. GraphPad Prism version 10 was employed to illustrate the growth inhibition.

### Anticlostridial activity of the potent hits against *C. difficile*

The microbroth dilution technique was used to test the antiviral hits against *C. difficile* ATCC BAA 1870 to identify the minimum inhibitory concentration (MIC) of the promising hits [24–26]. To obtain a final bacterial concentration of 5 x 10^5^ CFU/ml in 96-well plates, the hits and the control antibiotic, vancomycin, were serially diluted to a bacterial suspension of 0.5 McFarland solution in BHIS broth. These plates were then incubated anaerobically at 37°C for 48. The lowest concentration at which a substance could totally prevent bacterial growth was identified as the minimum inhibitory concentration, or MIC. Based on the MIC data of the active hits (MIC range ≤ 4μg/mL), four antiviral drugs (rottlerin, dryocrassin ABBA, α-mangostin, and obefazimod) were further studied. These four substances were purchased commercially and examined against a panel of fifteen clinical isolates of *C. difficile* in comparison to the control antibiotic: vancomycin. The test agent concentrations that inhibited 50% and 90% of the strains, respectively, were found to be the MIC_50_ and MIC_90_ values.

### Antibacterial activity of potent hits against representative members of gut microbiota

The most potent hits’ MICs against gut microbiota strains was ascertained in accordance with previous studies [27–29]. To achieve a final bacterial concentration of 5 x 10^5^ CFU/ml, a 0.5 McFarland bacterial solution was made in PBS and diluted for strains of *Bifidobacterium* and *Bacteroides* in BHIS broth and MRS broth for *Lactobacillus*. After serially diluting this suspension with the hits and control antibiotics, it was incubated for 48 hours at 37°C in anaerobic conditions for *Bifidobacterium* and *Bacteroides* or in the presence of 5% CO2 for *Lactobacillus* before determining the MICs of the hits, the lowest concentration at which it could inhibit completely bacterial growth.

### Time-kill kinetics assay against *C. difficile*

In a time-kill experiment, as previously described, potential antiviral drugs were challenged against *C. difficile* to ascertain the bactericidal or bacteriostatic nature of inhibition [20, 30]. *C. difficile* ATCC-43255 and *C. difficile* ATCC-BAA 1870 were grown overnight, then were diluted 1:150 into sterile BHIS yielding a concentration of approximately 5 × 10^5^ CFU/mL. *C. difficile* was treated with the hits and the control antibiotic, vancomycin, at a concentration of 5x MIC, and then cultured anaerobically at 37°C. As an untreated control, DMSO at a volume equal was also added. At 0, 2, 4, 8, 12, 24, and 48 hours, aliquots were collected from each treatment, diluted, and plated onto pre-reduced BHIS agar plates. Plates were incubated in anaerobic conditions at 37°C until the bacterial CFU could be counted. A drug was deemed to be bactericidal if it reduced the initial inoculum by ≥3-log10 CFU/mL.

## Results

### Screening and cherry-picking of the antiviral library against *C. difficile*

618 compounds from the MCE antiviral library were tested for possible anti-*C. difficile* action against the hypervirulent strain of *C. difficile* ATCC-BAA 1870. The screening was done at an initial concentration of 16 µM using the broth microdilution method. 24 compounds were identified with anticlostridial activity that inhibited the growth of *C. difficile* at the screening concentration (**Fig. 1**, **Table 1, and S1**). 15 hits that have previously been reported or identified as toxic or anticancer drugs were excluded based on findings from the preceding literature (**S1 Table**). The 9 novel hits had their activity confirmed against *C. difficile,* and their MICs were determined. Out of these 9 hits, four antiviral compounds (rottlerin, α-mangostin, dryocrassin ABBA, and obefazimod) displayed the most potent activity against *C. difficile* ATCC BAA-1870 with MIC values ranging from ≤4 μM (**Fig. 1**). Due to their unique nature and lack of prior research, these promising hits were chosen for further investigation. Among the most promising hits in the library (**Table 1**), 3 were natural products and 1 was a drug currently in clinical trials for HIV and ulcerative colitis treatments [31, 32]

**Figure 1.**
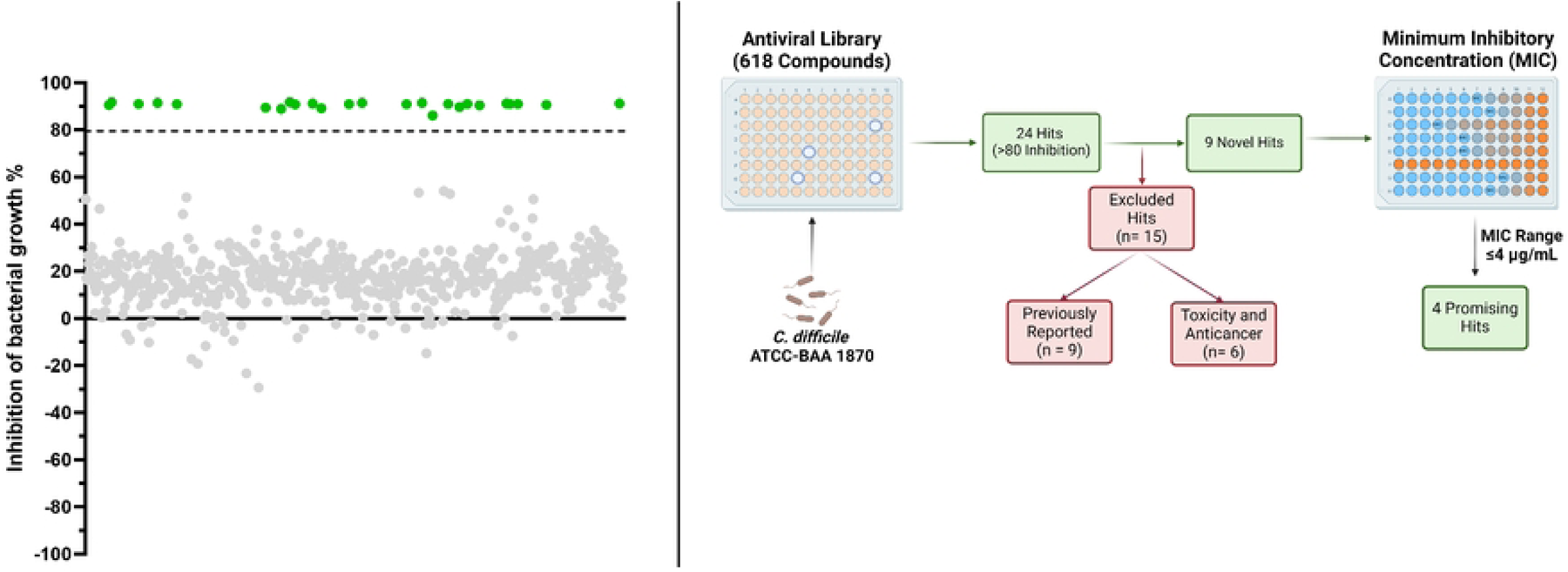
MCE antiviral library screening of 618 antivirals at 16 µM against *C. difficle* ATCC-BAA 1870. Compounds with a greater than 80% inhibition of bacterial growth were identified as hits (highlighted in green).

**Table 1.**
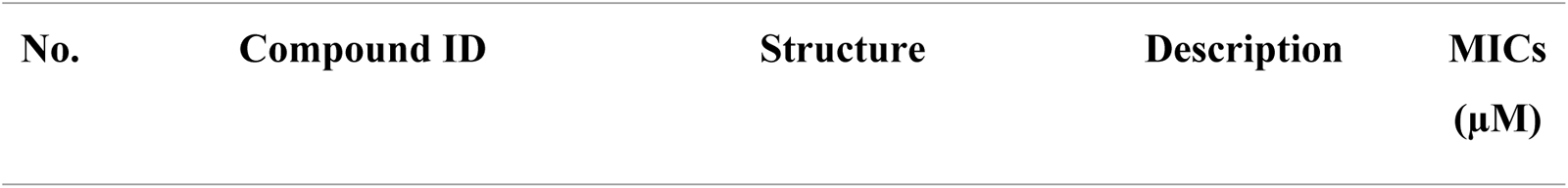
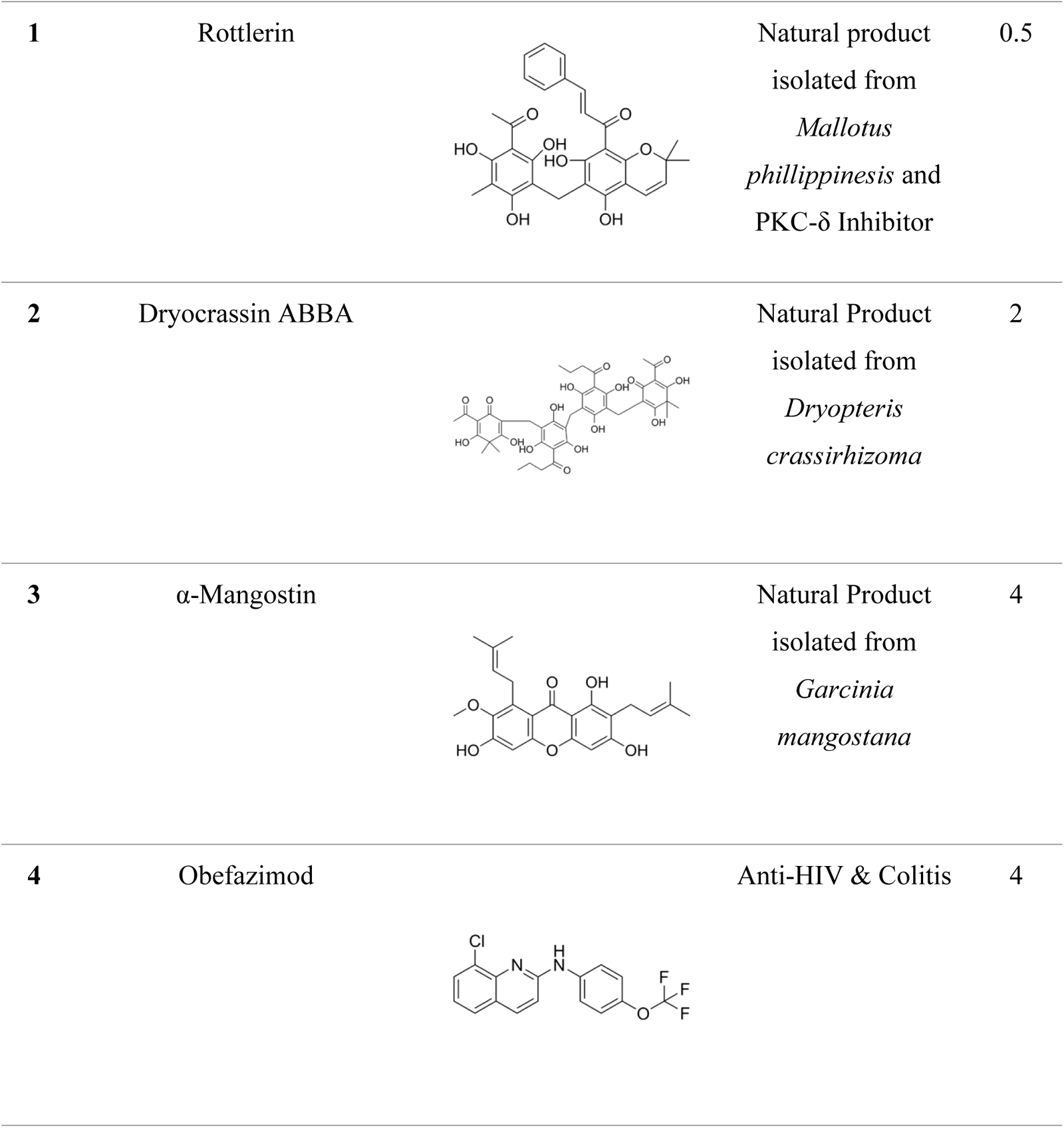
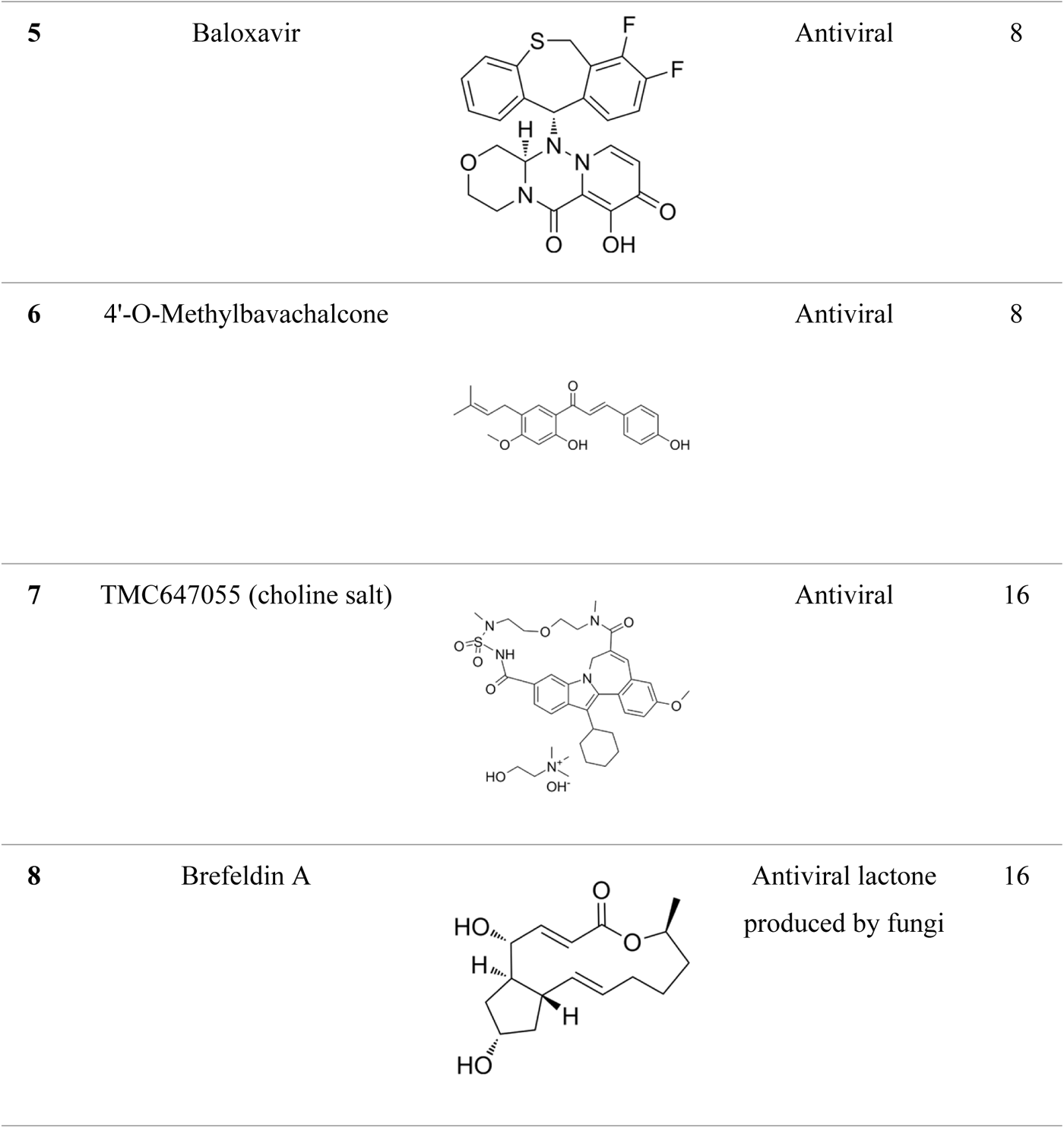
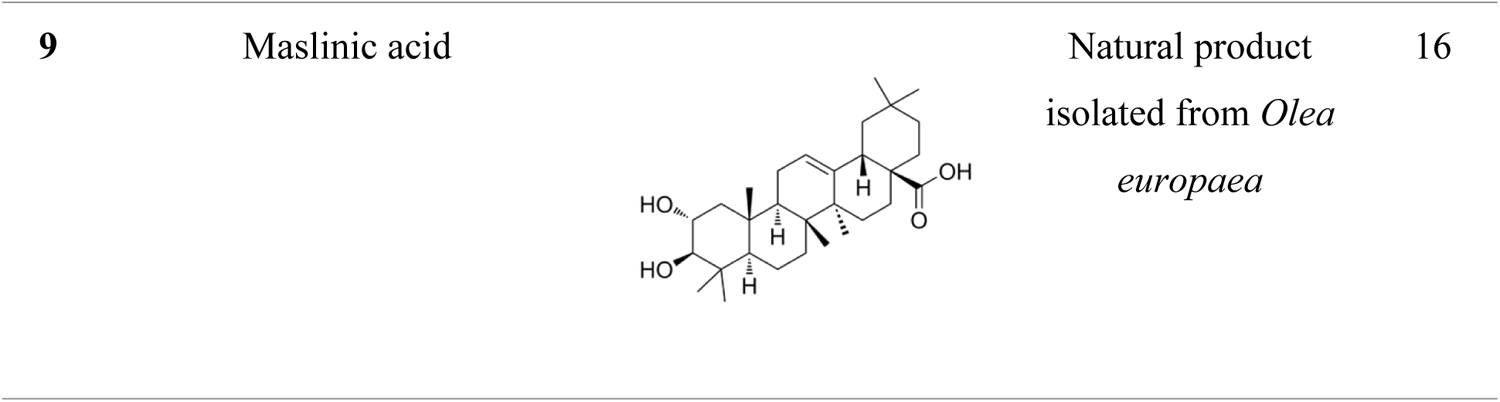
Minimum inhibitory concentrations (MICs) of potent antivirals against *C. difficile* ATCC BAA-1870. MICs are in µM concentration and are listed along with compound ID, chemical structure, and a brief description of their origin and/or function.

### Anticlostridial activity of the potent hits against *C. difficile*

The anticlostridial activity of the four top hits was further confirmed by testing them against 15 clinical isolates of *C. difficile* (**Table 2**) and calculating the MIC_50_ and MIC_90_ of each drug. Against the tested strains of *C. difficile*, the MIC values of rottlerin (MIC_50_ = 0.25 μg/mL and MIC_90_ = 0.5 μg/mL) were (MIC_50_ = 0.03 μg/mL and MIC_90_ = 0.06 μg/mL), whereas α-mangostin (MIC_50_ = 1 μg/mL and MIC_90_ = 2 μg/mL) was comparable to those of vancomycin (MIC_50_ = 0.5 μg/mL and MIC_90_ = 1 μg/mL). Likewise, dryocrassin ABBA (MIC_90_ = 4 μg/mL and MIC_50_ = 1 μg/mL) had similar activity to Vancomycin with a one-fold difference in its MIC_90_. Conversely, obefazimod needed a 3-to-4-fold greater concentration to inhibit *C. difficile* (MIC_50_ = 4 μg/mL and MIC_90_ = 8 μg/mL) in comparison to Vancomycin.

**Table 2.** The minimum inhibitory concentration (MIC) values of potent antivirals and antibiotics against clinical isolates of *C. difficile*. Against 15 clinical isolates of *C. difficile* with drug of choice vancomycin serving as a control.

| *C. difficile* strain ID | Minimum inhibitory concentration (µg/mL) |  |  |  |  |
| --- | --- | --- | --- | --- | --- |
|  | Antiviral hits |  |  |  | Control antibiotic |
| | Rottlerin | α-Mangostin | Dryocrassin ABBA | Obefazimod | Vancomycin |
| ATCC-BAA 1870 | 0.5 | 1 | 2 | 4 | 1 |
| ATCC 1871 | 0.5 | 2 | 4 | 8 | 2 |
| ATCC 43255 | 0.25 | 0.5 | 0.5 | 2 | 0.5 |
| ATCC 43598 | 0.5 | 1 | 1 | 8 | 0.5 |
| BAA 630 | 0.25 | 0.5 | 1 | 4 | 0.5 |
| NR-49288 | 0.5 | 2 | 4 | 8 | 1 |
| NR-49302 | 0.5 | 0.5 | 2 | 4 | 0.5 |
| NR-49308 | 0.25 | 0.5 | 0.5 | 4 | 0.5 |
| NR-49313 | 0.25 | 1 | 0.5 | 2 | 0.5 |
| NR-49319 | 0.5 | 1 | 1 | 4 | 0.25 |
| CDI-12 | 0.25 | 1 | 0.5 | 4 | 1 |
| CDI-16 | 0.25 | 0.5 | 1 | 2 | 0.5 |
| CDI-20 | 0.25 | 0.5 | 1 | 4 | 0.5 |
| CDI-21 | 0.5 | 2 | 4 | 8 | 1 |
| CDI-28 | 0.25 | 1 | 1 | 8 | 1 |
| MIC<sub>50</sub> | **0.25** | **1** | **1** | **4** | **0.5** |
| MIC<sub>90</sub> | **0.5** | **2** | **4** | **8** | **1** |

### Antibacterial activity of the potent hits against representative members of gut microbiota

To test the activity against gut microflora, selected antiviral compounds were screened against a panel of representative commensal gut microbiota *Lactobacillus*, *Bacteroides*, and *Bifidobacterium* up starting at a concentration of 256 µg/mL. The selected hits and vancomycin had little to no effect on *Lactobacillus* strains apart from fidaxomicin (MIC values of 8 and 32 µg/mL against *rhamnosus* and *brevis* strains, respectively) (**Table 3**). Rottlerin, obefazimod, and dryocrassin ABBA showed activity at higher concentrations (MIC values ranged from 8 to 128 µg/mL) against *Bacteroides fragilis* strains in comparison to vancomycin (MIC values, 16-32 µg/mL), however, α-mangostin displayed potent activity at lower concentrations (MIC values, ≤2 µg/mL), while fidaxomicin showed no activity at the maximum tested concentration (MIC values, >256 µg/mL) (**Table 3**). Rottlerin, obefazimod, and dryocrassin ABBA show activity at lower concentrations (MIC values, 4-8 µg/mL) against *Bifidobacterium breve* strains, whilst α-mangostin shows activity at lower concentrations (MIC values, ≤2 µg/mL) similar to the control antibiotics, vancomycin and fidaxomicin (**Table 3**). Overall, antiviral compounds showed similar or less activity against gut microflora when compared to drugs of choice for CDI with the greatest difference in activity being against *Bifidobacterium* strains being at least 2-4 folds less active.

**Table 3.**
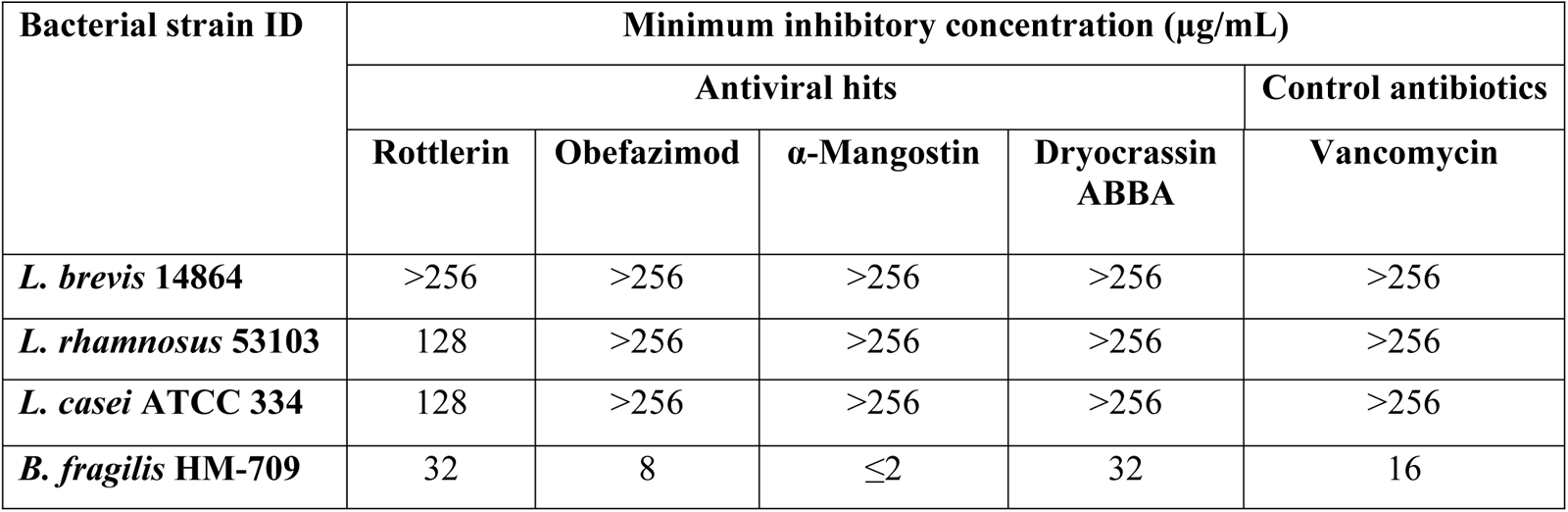
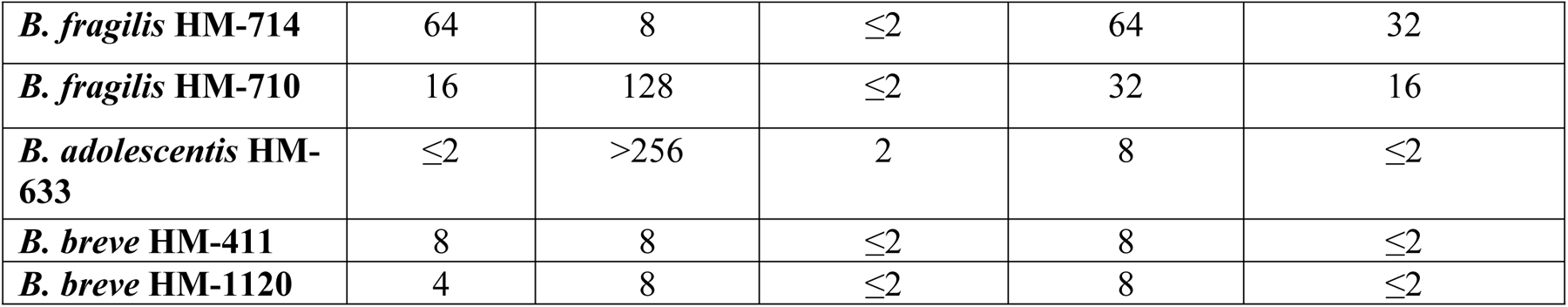
MIC (µg/mL) values of selected antivirals and control antibiotics against representative members of human normal gut microbiota.

| Bacterial strain ID | Minimum inhibitory concentration (µg/mL) |  |  |  |  |
| --- | --- | --- | --- | --- | --- |
|  | Antiviral hits |  |  |  | Control antibiotics |
| | Rottlerin | Obefazimod | α-Mangostin | Dryocrassin<br>ABBA | Vancomycin |
| *L. brevis* 14864 | >256 | >256 | >256 | >256 | >256 |
| *L. rhamnosus* 53103 | 128 | >256 | >256 | >256 | >256 |
| *L. casei* ATCC 334 | 128 | >256 | >256 | >256 | >256 |
| *B. fragilis* HM-709 | 32 | 8 | $\leq 2$ | 32 | 16 |
| *B. fragilis* HM-714 | 64 | 8 | $\leq 2$ | 64 | 32 |
| *B. fragilis* HM-710 | 16 | 128 | $\leq 2$ | 32 | 16 |
| *B. adolescentis* HM-633 | $\leq 2$ | >256 | 2 | 8 | $\leq 2$ |
| *B. breve* HM-411 | 8 | 8 | $\leq 2$ | 8 | $\leq 2$ |
| *B. breve* HM-1120 | 4 | 8 | $\leq 2$ | 8 | $\leq 2$ |

### Time-kill kinetics assay against *C. difficile*

To assess the killing kinetics (bactericidal or bacteriostatic) activity of the antiviral compounds, a time-kill assay was performed against *C. difficile* ATCC-BAA 1870 and ATCC 43255. As demonstrated in **Fig. 3A, B**, α-mangostin exhibited strong bactericidal activity within 2 hours, significantly better than the control antibiotics Fidaxomicin and vancomycin, which demonstrated bactericidal activity at 12 hours and 24 hours, respectively, for both strains. In contrast, dryocrassin ABBA, rottlerin, and obefazimod displayed bacteriostatic activity with approximately 1-1.5 log reduction within 12 hours and remained static up to 48 hours (**Fig. 3A, B**).

**Figure 2.**
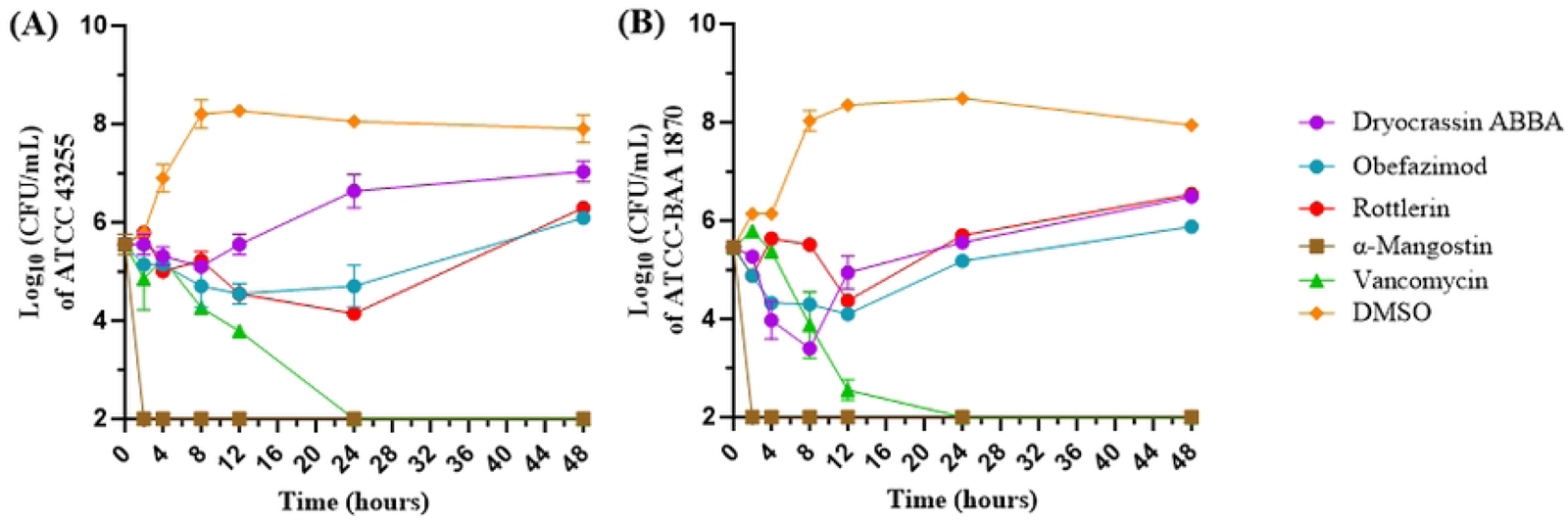
Time-kill assay of the potent hits and the control antibiotics against *C. difficile* ATCC 43255 (A) or ATCC-BAA 1870 (B) at 5x MIC. Performed on pre-reduced BHIS medium. *C. difficile* development was observed as colony forming units over a 24-hour period.

## Discussion

*C. difficile* is one of the most reported and impactful nosocomial pathogens in the United States, accounting for $1.5 billion in annual expenditures [7]. The issue of the CDI recurrence and potential for the development of antimicrobial resistance is still present in the two drugs of choice for CDI [33] [13]. This highlights a growing need for a novel, more consistent, and effective drug or drug scaffold to combat CDI. Antibiotics for associated secondary infections have disrupted the gut microbiota, increasing the likelihood of *C. difficile* colonization, which has made CDI a worry for patients receiving treatment for viral infections[34, 35]. Thus, discovering antivirals with strong anti-*C. difficile* action is a fruitful approach that clinics prioritize highly to lower the risk of CDI.

Therefore, in this study, a whole cell-based screening of a library of antiviral compounds was conducted against *C. difficile*. In the initial screening, 24 compounds were found to have anti-*C. difficile* activity. After excluding anticancer compounds and previously reported hits, we pursued 9 antivirals (**Table 1**). These results were confirmed and refined, ultimately narrowing down to 4 compounds that displayed the most potent activity against *C. difficile* ATCC BAA-1870 with MIC values ≤4 μM. Due to their unique nature and lack of prior research regarding their use as antimicrobials, these promising hits were chosen for further investigation. The 4 compounds identified were rottlerin, α-mangostin, dryocrassin ABBA, and obefazimod. To further confirm their activity, these hits were purchased commercially and put against a panel of 15 clinical isolates of *C. difficile* to determine their MIC_50_ and MIC_90_ (**Table 2**).

Rottlerin, also known as mallotoxin, is a natural product isolated from *Mallotus phillippinesis* and is a known protein kinase Cδ (PKCδ)-selective inhibitor and mitochondrial uncoupler that is typically utilized as the base for substrate phosphorylation studies [36, 37]. However there has been some debate on whether it is selective for protein kinase C [37]. Rottlerin has displayed activity targeting quorum sensing and biofilm activity in *Pseudomonas aeruginosa* [38]. It has also shown activity against *mycobacteria spp.*, including *M. tuberculosis* and *M. smegmatis* (IC_50_ range 9-74 µM), potentially interfering with shikimate kinase, a promising target for antimicrobials, which is essential for many bacteria [39]. It also has been reported to target *Chlamydia* species with MIC values of 1 µM [40]. Recently, rottlerin was found to have some fungicidal activity in a study where MIC and MFC values against *Candida* species with values as low as 7.81 µg/mL [41]. Here, we report that rottlerin exhibited strong antibacterial activity against *C. difficile* (MIC_50_= 0.25μg/mL and MIC_90_= 0.5μg/mL), which was comparable to the control antibiotic, vancomycin.

α-mangostin is a xanthone derivate extracted from the edible fruit of *Garcinia mangostana*, commonly known as mangosteen [42]. α-mangostin showed potent anti-*C. difficile* activity (MIC_50_= 1μg/mL and MIC_90_= 2μg/mL) (**Table 2**). Interestingly, α-mangostin has faced issues as a treatment due to its lack of absorption and oral bioavailability, leading researchers to modify α-mangostin to improve its pharmacokinetics [43]. Interestingly, these characteristics are a benefit for CDI treatment which make it an intriguing molecule to further pursue for development as a novel anti-*C. difficile* agent. Both primary treatments for moderate to severe CDI, vancomycin and fidaxomicin, have minimal systemic absorption when taken orally [44, 45]. α-mangostin has been found in high concentrations within the small intestine, making it ideal for treatment of *C. difficile*. In addition, α-mangostin has shown potent and rapidly bactericidal activity both in this study and against methicillin-resistant *Staphylococcus aures* (MRSA) [46–48]. This compound was potent in *C. difficile* equivalent to that of drug of choice vancomycin (**Table 2**) with little to no activity against microbiota (Table 3) and a far more rapid mechanism of killing (Figure 2).

Dryocrassin ABBA is a phloroglucinol derivative that is isolated from *Dryopteris crassirhizoma* [49] and has previously been reported to have antimicrobial activity against both H5N1v avian influenza virus and *Staphylococcus aureus* (*S. aureus)* virulence factors [50, 51]. Particularly, it has potent activity against virulence factors of MRSA such as Von Willebrand factor-binding proteins (vWbp) and Sortase A [51, 52]. Otherwise, it does not have activity against *S. aureus* itself (MIC >1024 µg/mL) [51]. To contrast, dryocrassin ABBA showed activity similar to that of vancomycin against *C. difficile*, with an MIC_50_ and MIC_90_ of 1 µg/ml and 4 µg/ml (**Table 1**).

Obefazimod, also known as ABX464, is a novel and first-in-class compound developed by Abivax and has shown antiviral, antirheumatic, and anti-colitis properties [31, 53, 54]. It is currently in phase 2 trials for its antiviral and anti-rheumatic properties, while in phase 3 trials for anti-colitis effect [31, 32, 55, 56]. There have been no previous reports of antibacterial activity published so far, making this compound of great interest as a potential novel class of antimicrobials on top of its other therapeutic effects. This compound had an MIC_50_ of 4 µg/mL and MIC_90_ of 8 µg/mL (**Table 2**) and displayed bacteriostatic activity against *C. difficile* (Figure 2). Its activity against human microbiota was lower across the board when compared to control antibiotics apart from *Bacteroides* strains (**Table 3**).

*C. difficile* is an opportunistic pathogen that is intimately tied with gut dysbiosis. Administration of broad-spectrum antibiotics can disrupt commensal gut microflora and remove the first line of defense against *C. difficile* colonization [57]. Due to this, selectivity against *C. difficile* could be beneficial in preventing recurrence and reduce the severity of CDI[58]. Therefore, it is necessary to take into consideration whether these compounds have an impact on beneficial bacteria typically found in the human gut microbiome. Overall, the selected compounds had very little effect on gut microbiota and in some cases had several folds MICs higher than that of drugs of choice. For *Bifidobacterium* strains of bacteria, all compounds with the exception α-mangostin had MICs several fold higher than both drugs of choice, vancomycin and fidaxomicin.

Next, we aimed to monitor the killing kinetics of the selected antiviral compounds to determine whether they had bactericidal or bacteriostatic activity. We found that 3 compounds, rottlerin, dryocrassin ABBA, and obefazimod had bacteriostatic activity. However, α-mangostin displayed rapid bactericidal far more potent than both fidaxomicin and vancomycin, achieving 100% inhibition within the first 2 hours. Remarkably, α-mangostin’s potent activity against *C. difficile* (MIC_50_= 1μg/mL and MIC_90_= 2μg/mL) (**Table 2**) did not translate to other bacteria with little to no activity against microbiota (**Table 3**) and a far more rapid mechanism of killing (**Fig. 2**) highlighted the potential of this compound for CDI treatment.

To conclude, the goal of this work was to screen antiviral libraries for new anti-*C. difficile* compounds, such as obefazimod, rottlerin, α-mangostin, and dryocrassin ABBA, that showed strong anti-*C. difficile* activity. Most notably, α-mangostin’s killing kinetics were superior to those of the preferred medications now. When it comes to avoiding recurrent CDI, commensal microbiota is highly advantageous, and nearly all drugs shown little to no efficacy against it. The compounds found in this screening could serve as lead structures for further development.

## Competing interests

None declared.

## Acknowledgements

The authors would like to thank BEI Resources and the CDC for providing some of the strains used in this study. This work was funded by the National Institutes of Health (grant R01AI130186).

## Notes

### Competing Interest Statement

The authors have declared no competing interest.

